# Body schema plasticity is altered in Developmental Coordination Disorder

**DOI:** 10.1101/2021.08.16.456453

**Authors:** Marie Martel, Véronique Boulenger, Eric Koun, Livio Finos, Alessandro Farnè, Alice Catherine Roy

## Abstract

Developmental Coordination Disorder (DCD) is a pathological condition characterized by impaired motor skills. Current theories advance that a deficit of the internal models is mainly responsible for DCD children’s altered behavior. Yet, accurate movement execution requires not only correct movement planning, but also integration of sensory feedback into body representation for action (Body Schema) to update the state of the body. Here we advance and test the hypothesis that the plasticity of this body representation is altered in DCD. To probe Body Schema (BS) plasticity, we submitted a well-established tool-use paradigm to seventeen DCD children, required to reach for an object with their hand before and after tool use, and compared their movement kinematics to that of a control group of Typically Developing (TD) peers. We also asked both groups to provide explicit estimates of their arm length to probe plasticity of their Body Image (BI). Results revealed that DCD children explicitly judged their arm shorter after tool use, showing changes in their BI comparable to their TD peers. Unlike them, though, DCD did not update their implicit BS estimate: kinematics showed that tool use affected their peak amplitudes, but not their latencies. Remarkably, the kinematics of tool use showed that the motor control of the tool was comparable between groups, both improving with practice, confirming that motor learning abilities are preserved in DCD. This study thus brings evidence in favor of an alternative theoretical account of the DCD etiology. Our findings point to a deficit in the plasticity of the body representation used to plan and execute movements. Though not mutually exclusive, this widens the theoretical perspective under which DCD should be considered: DCD may not be limited to a problem affecting the internal models and their motor functions, but may concern the state of the effector they have to use.

## Introduction

Developmental Coordination Disorder (DCD) is a neurodevelopmental condition marked by impaired motor skills in the absence of neurological injury, given a child’s chronological age and previous opportunities for skill acquisition (DSM-V: American Psychiatric Association, 2013). It affects approximately 5% of school-aged children (Mandich and Polatajko, 2003; range between 1.8% and 8% depending on the selection criteria; for reviews see Biotteau et al., 2020; Gomez and Sirigu, 2015; Zwicker et al., 2012), and persists through adulthood (e.g. Cantell et al., 2003; Cousins and Smyth, 2003; Losse et al., 1991; Rasmussen and Gillberg, 2000), thus considerably impacting academic and life achievements (e.g. Cheng et al., 2011; Geuze, 2005; Kirby and Sugden, 2007; Smits-Engelsman et al., 2001; Van der Linde et al., 2015).

The etiology of DCD has gained considerable interest over the years, with converging evidence towards an internal modelling deficit (also called predictive control; Adams et al., 2014; Gomez and Sirigu, 2015; Wilson et al., 2013). Within internal models, the inverse model defines the accurate motor command, and the forward model predicts its consequences. Actual sensory feedback are also monitored allowing for movement correction and update of both models to improve motor control (Kawato and Wolpert, 1998; Shadmehr and Krakauer, 2008). Children with DCD can produce some appropriate motor commands when required, even though these appear to be more variable than in typically developing (TD) children (e.g. Roche et al., 2016; Smits-Engelsman et al., 2008; Smits-Engelsman and Wilson, 2013). Accordingly, visuomotor adaptation studies showed that children with DCD are able to update their inverse model, provided they are given more trials and larger error signals (Cantin et al., 2007; Kagerer et al., 2006, 2004; Zoia et al., 2005). This suggests that they need more time to process feedback, and that they might ignore error signals that are not relevant enough. When relevant, however, they can learn from these signals and update their models, as shown by their preserved ability for motor learning (Smits-Engelsman et al., 2015). Furthermore, children with DCD plan and execute a whole movement (i.e., in one time), rather than performing an incomplete movement and updating it online, as this may seem too costly for their motor system (Mon-williams et al., 2005). Hence, in double-step paradigms, children with DCD show difficulties in correcting 3D movement trajectories through rapid online control (Hyde and Wilson, 2013, 2011a, 2011b). Yet, they have no deficit when tasks are easier, namely when more time is allowed to correct the movement, or in tasks that involve 2D movements in the transversal plane (Adams et al., 2016; Plumb et al., 2008). Online control performance can be predicted by the ability to represent action, that is motor imagery abilities, both in typical and atypical development (Fuelscher et al., 2015a, 2015b). Motor imagery studies, which tackle the integrity of the forward model (Kilteni et al., 2018), revealed that DCD participants can imagine movements, but again less consistently, less accurately and less rapidly than their TD peers (e.g. Barhoun et al., 2021, 2019; Deconinck et al., 2009; Noten et al., 2014; Reynolds et al., 2015; Williams et al., 2008; Wilson et al., 2004). Children with DCD can make proper use of instructions to improve their motor imagery abilities (Reynolds et al., 2015; Williams et al., 2008), and motor imagery training has been shown helpful for motor control remediation in DCD (Steenbergen et al., 2020), again attesting that internal models can be updated. Overall, deficits in DCD therefore seem to affect different components of internal models, making it difficult to decipher the core neurocognitive deficit.

A so far relatively neglected aspect of the internal models is the body estimate. This body representation for action, also referred to as Body Schema (BS), is defined as an implicit (i.e., unconscious), plastic sensorimotor representation that allows monitoring the position and size of the different effectors (de Vignemont, 2010; Head and Holmes, 1911; Martel et al., 2016; Medina and Coslett, 2010; Schwoebel and Coslett, 2005). This representation is often opposed to a body representation for perception, the so-called Body Image (BI), which is an action-free, explicit (i.e., conscious) representation of the body shape and size (de Vignemont, 2010; Gallagher, 2005). Owing to its action-devoted function, the BS (but not the BI) is referred to as body state estimation in the motor control domain. Here, we will therefore refer to each specific body representation accordingly. Substantial evidence for the plastic monitoring of limb’s size in motor control comes from studies on tool use. After using a tool for a few minutes, healthy adults start performing free-hand movements differently, with longer latencies and reduced amplitudes for the acceleration, velocity and deceleration profiles (e.g. Cardinali et al., 2009); for review Martel et al., 2016). This kinematic signature of what has been called tool-incorporation, typically observed also in long-armed vs. short-armed participants (Cardinali et al., 2012; Martel et al., 2019), is indicative of a longer arm estimate after tool use. This body state plasticity has been suggested to allow tools to become extensions of our limbs for action and perception (e.g. Arbib et al., 2009; Jacobs et al., 2009; Maravita and Iriki, 2004; Martel et al., 2016; Miller et al., 2019, 2018; Witt, 2021).

Importantly, this plasticity requires years to develop fully. Martel and colleagues (2021) reported that the plasticity of the state estimation is not mature in TD children and early adolescents. After using a tool for a few minutes, their free-hand kinematic pattern is actually opposite to what has been consistently observed in healthy adults: the amplitude peaks of the wrist increased, and their latencies decreased. Following the rationale from previous studies in adults, this was interpreted as a movement performed with an arm estimated as being shorter (rather than longer in adults) after use of the same tool. Reduction in arm length estimate may result from the need to build new sensorimotor associations for a tool that has never been used before (Ganesh et al., 2014) or from stronger reliance on visual as compared to proprioceptive guidance in TD children, compared to adults (Martel et al., 2021). Investigating the plasticity of the body estimate in DCD may offer complementary insights to better understand its aetiology (Martel et al., 2021, 2019, 2017; Medendorp and Heed, 2019).

Surprisingly, the potential role played by an altered body estimate in DCD’s motor deficits has yet been scarcely investigated (Gomez and Sirigu, 2015), although the two have been linked indirectly. Impairment in multisensory body representations have been suggested after findings of poorer performance in somatosensory localization in DCD participants (Elbasan et al., 2012; Johnston et al., 2017; Schoemaker et al., 2001). Wilson and colleagues (2004) also hypothesized that motor imagery difficulties in DCD could result from inaccurate body estimate: children with DCD would not automatically use motor imagery from their perspective, but would rather rely on visual imagery from an object-based third-person perspective (see also Barhoun et al., 2021).

Here, we aimed at filling this gap by investigating the plasticity of both the implicit body estimate (Body Schema) and the explicit Body Image in DCD, taking advantage of the above-mentioned tool-use paradigm. First, given their relatively preserved ability to update the inverse model and program movements (e.g. Kagerer et al., 2006; Smits-Engelsman et al., 2015, 2008), we predicted that children and early adolescents with DCD should be able to control the tool adequately and to improve with practice. Quantitative difference with their TD peers might however be expected as they sometimes ignore sensory signals, or need more trials to reach the same level. Second and most importantly, we predicted that children with DCD would show a less plastic body estimate than their TD peers. Accordingly, free-hand movements following tool use should display a different kinematic signature than that observed in TD children. Lastly, we also investigated the conscious, explicit representation (or Body Image) in DCD and TD, asking them to estimate their perceived forearm length before and after tool use. With respect to the Body Image and its plasticity, we anticipated that it would be preserved. Indeed, the BI is not linked to motor control, making any disruption in DCD unlikely. Moreover, clinical assessment of the ability to point or name several body-parts is generally preserved in DCD children, suggesting they can access their perceptual body metrics. Thus, their consciously perceived forearm length should be judged as shorter after tool use, as recently observed in TD children (Martel et al., 2021).

## Material and methods

### Participants

Owing to the limited number of developmental studies involving tool-use, particularly in DCD children (Caçola et al., 2014; Caçola and Gabbard, 2012), we recruited participants during a full school year (9 months, 2015-2016), in parallel to the recruitment of TD participants from a previous study using the same paradigm (Martel et al., 2021). A total of 32 DCD participants, aged between 9.5 and 16.5 years old, were referred to our lab by several physical therapists as well as by the health network Dys/10 (specialized in the care of developmental disorders, from diagnosis with a multidisciplinary team to follow-ups with schools). All children were diagnosed with DCD following the DSM-5 (American Psychiatric Association, 2013), and accordingly had no neurological condition nor intellectual disability. They were additionally screened on the French adaptation of the Movement ABC (Henderson and Sugden, 1992; Soppelsa and Albaret, 2004) by a trained physical therapist (duration ~ 1h30). As the French version of the M-ABC2 (Marquet Doléac et al., 2016), standardized for adolescents up to 16 years of age, was not available at the time of the study, all the adolescents with DCD older than 12 years were evaluated based on the standardization for the 12-year-olds. To be included in the final sample, participants with DCD could not be born preterm (n = 2 excluded) and had to score below the 5^th^ percentile on the M-ABC, or below the 15^th^ percentile but with at least one subcategory under the 5^th^ percentile (n = 5 excluded). Comorbidity with other developmental disorders was not an exclusion criterion (see Table 1).

**Table 1.**
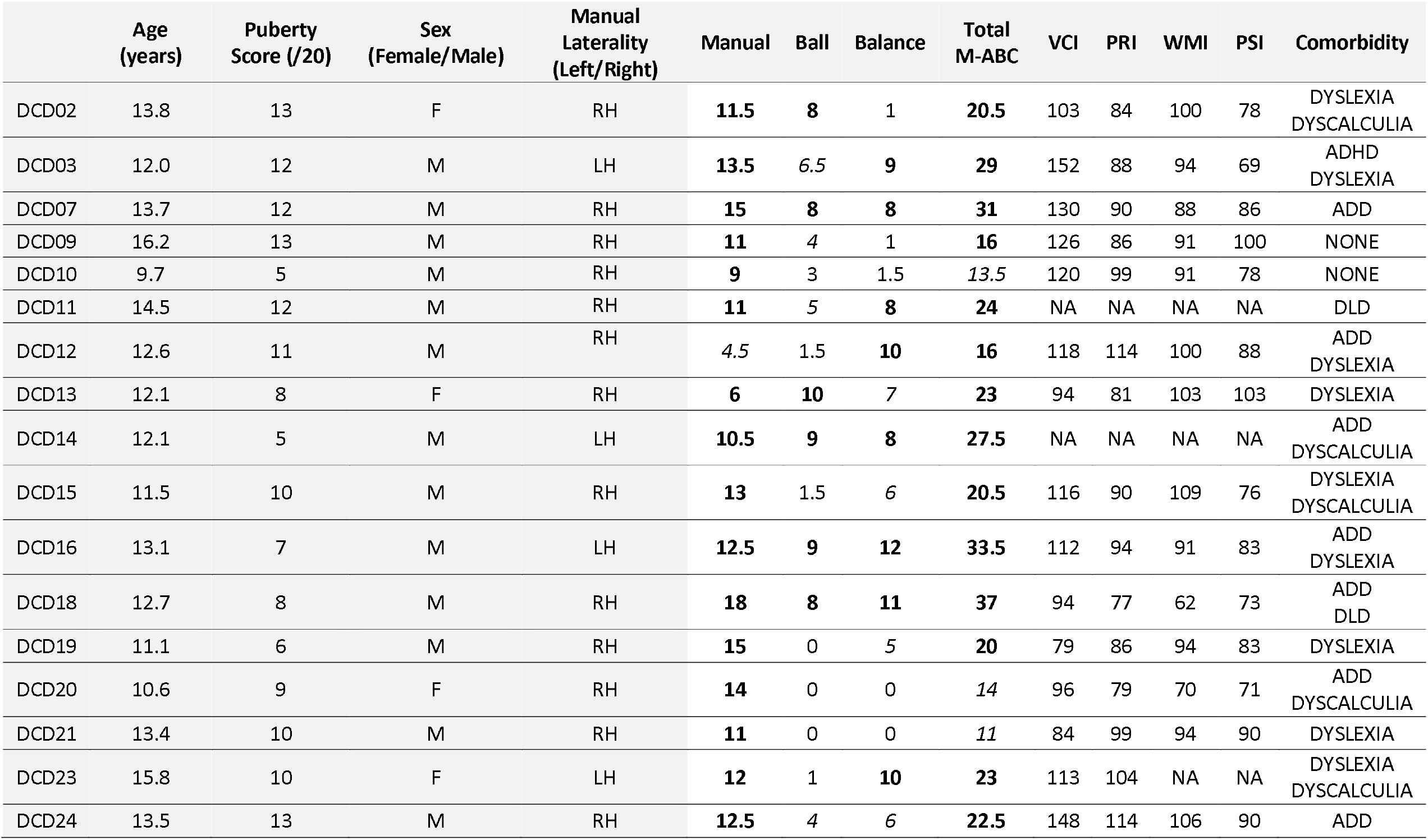
Characteristics of the group of children and early adolescents with DCD (n=17). The scores on manual dexterity, ball skill and balance are the three subtests of the M-ABC. The scores ≤ 5^th^ percentile are indicated in bold, and those ≤ 15^th^ percentile in italics. Scores for the VCI (Verbal Comprehension Index), PRI (Perceptual Reasoning Index), WMI (Working Memory Index) and PSI (Processing Speed Index) are subtests of the IQ test (Grizzle, 2011). NA indicates missing data for participants referred by physical therapists but whose parents could not provide us with any IQ assessment score. Comorbidity with other developmental disorders such as dyslexia, dyscalculia, Developmental Language Disorder (DLD) or Attention Deficit Disorder with (ADHD) or without Hyperactivity (ADD) is indicated.

Out of the 25 remaining DCD participants who fulfilled these criteria, 17 could be included in the current study to meet the criterion of puberty level (see below). They were matched with a group of 17 TD participants included in another study conducted by our group at the same time (Martel et al., 2021), according to their puberty level, sex, age, and practice of tool-based activities. We deemed important to select our sample on the basis of puberty (instead of age) because puberty is known to impact sensorimotor development importantly with, for instance, altered use of proprioception (Assaiante et al., 2014; Barlaam et al., 2012; Cignetti et al., 2013; Viel et al., 2009), impoverished integrative performance (Nardini et al., 2013) and transient clumsiness linked to the growth spurt (Hirtz and Starosta, 2002; Martel et al., 2021; Visser and Geuze, 2000). It was thus crucial to ensure that participants would be at a comparable puberty level, irrespective of their age. We assessed the puberty level with the Self-Rating Scale for Pubertal Development (Carskadon and Acebo, 1993; Petersen et al., 1988) including five items standing for five phenomena: growth spurt, body hair, skin changes, deepening of the voice/breast growth and growth of hair on the face/menstruations. For each item, puberty-induced changes are rated on a 4-point scale (“not started yet”, “barely started”, “definitely underway”, “finished”), leading to a global puberty score (ranging between 5 and 20). This puberty score is an individual accurate estimation of body growth and indirectly accounts for the effect of sex/age. Adolescents who would indicate that each of the phenomena was “definitely underway” would score 15/20, meaning that they reached their puberty peak, while if all of the phenomena had “barely started”, they would score 10/20. We placed the cut-off for inclusion at 13/20 based on the results in typically developing children and adolescents using the same paradigm (Martel et al., 2021), which showed a change in tool-use effects on their body representations at mid-puberty (score ≥ 14), corresponding to the growth spurt. By including only children and early puberty adolescents with DCD scoring between 5 and 13/20, we minimized the potential confound of puberty, and focused on a period where the pattern of results in TD participants is rather stable. Importantly here, age was not relevant for our decision as children of the same age/sex have not necessarily reached the same pubertal stage and we fully relied on the puberty score. This led to a final sample of 17 DCD participants, who were compared to a subsample of 17 participants from a larger cohort of TD participants from a previous study (Martel et al., 2021). The two groups were matched for daily tool practice, age, sex and puberty (see Table 2). TD children and early adolescents had no learning disabilities or delayed psychomotor acquisition according to their parents’ report. DCD and TD participants all had normal or corrected-to-normal vision. We also ensured that DCD and TD groups did not differ in their sport and/or musical practice or in their daily tool-use activities (i.e. activities involving tools such as tennis, ping pong, golf, drums etc. that are regularly practiced outside school). Non-tool users practiced activities without any tool involved (e.g., football, handball, climbing, piano…). Matching DCD and TD participants for tool-use activities ensured that any difference in performance between the groups could not result from different knowledge in such tool-based activities. The main characteristics of the two groups are displayed in Table 2.

**Table 2.**
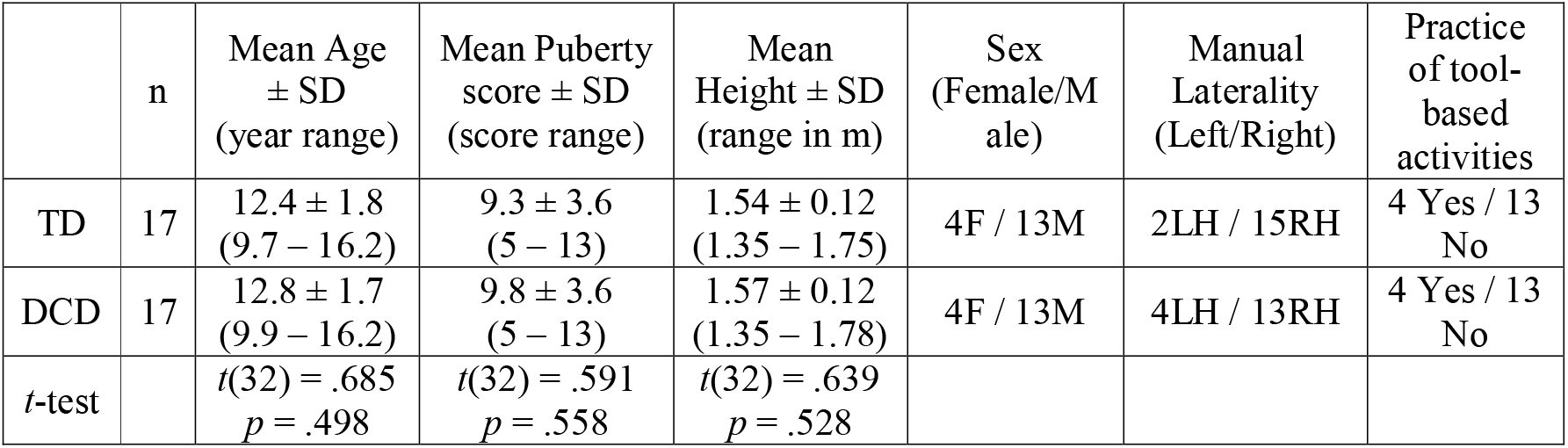
Characteristics of the Typically Developing and the Developmental Coordination Disorder groups.

|  | n | Mean Age<br>± SD<br>(year range) | Mean Puberty<br>score ± SD<br>(score range) | Mean<br>Height ± SD<br>(range in m) | Sex<br>(Female/M<br>ale) | Manual<br>Laterality<br>(Left/Right) | Practice<br>of tool-based<br>activities |
| --- | --- | --- | --- | --- | --- | --- | --- |
| TD | 17 | 12.4 ± 1.8<br>(9.7 – 16.2) | 9.3 ± 3.6<br>(5 – 13) | 1.54 ± 0.12<br>(1.35 – 1.75) | 4F / 13M | 2LH / 15RH | 4 Yes / 13<br>No |
| DCD | 17 | 12.8 ± 1.7<br>(9.9 – 16.2) | 9.8 ± 3.6<br>(5 – 13) | 1.57 ± 0.12<br>(1.35 – 1.78) | 4F / 13M | 4LH / 13RH | 4 Yes / 13<br>No |
| *t*-test |  | *t*(32) = .685<br>*p* = .498 | *t*(32) = .591<br>*p* = .558 | *t*(32) = .639<br>*p* = .528 |  |  |  |

Parents or guardians as well as children and early adolescents gave written informed consent to participate in the study, which was approved by the French ethics committee (*Comité de Protection des Personnes* CPP Sud-Est II) and conformed to the Helsinki declaration. Children and early adolescents were naïve to the purpose of the study and received a board game after completion.

### Apparatus and procedures

The paradigm was identical to the one used in a previous study in TD children and adolescents (Martel et al., 2021). This paradigm has consistently been shown to be sensitive to changes in arm length implicit estimate after tool use (see, for review, Martel et al., 2016). In summary, participants were required to reach-to-grasp an object with their free dominant hand before and after performing reach-to-grasp movements towards the same object with a tool lengthening their arm. Comparison of the kinematics of free-hand movements before and after tool use allows to measure the effects of tool use on free-hand movements, and thus to assess the plasticity of the body representation for action (Body Schema). We also assessed the effect of tool use on the subjective estimate of the forearm length (Body Image) by asking participants to estimate the length of their forearm before and after performing similar reach-to-grasp movements with the same tool. Figure 1 provides a schematic of the experimental design.

**Figure 1.**
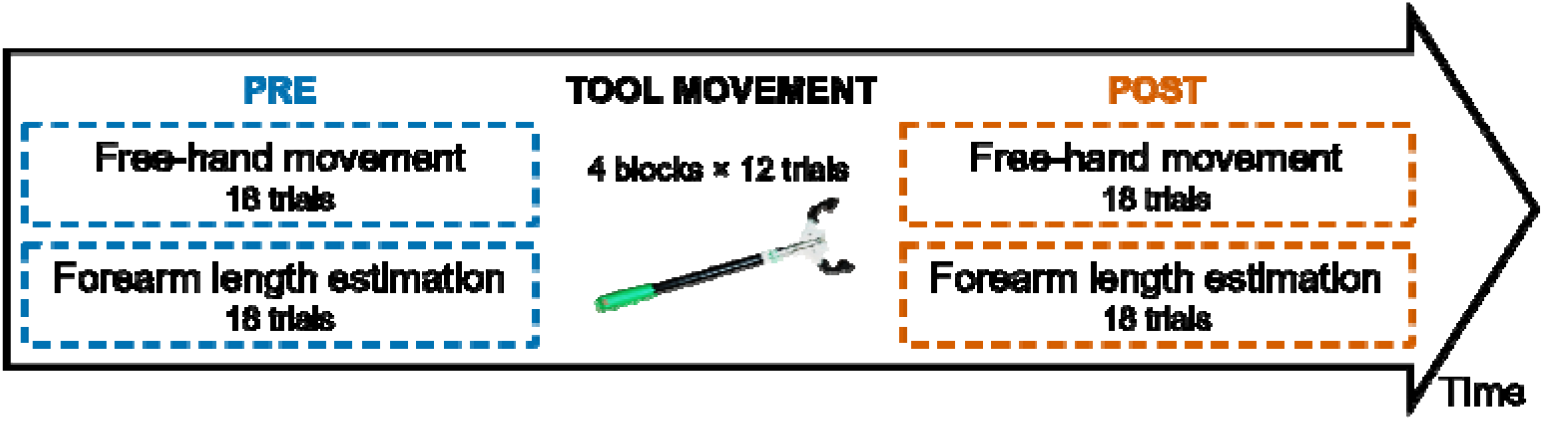
Experimental design of the tool-use paradigm. The two tasks in PRE and POST sessions were identical and counterbalanced between participants.

Participants were comfortably seated on an adjustable swivel chair, at a fixed distance from a table in front of them. The experiment was divided into three sessions: a pre- and post-tool-use session, separated by a tool-use session. During both pre- and post-tool-use sessions, participants performed a Free-Hand Movement task, and a Forearm Length Estimation task, counterbalanced across participants.

#### Free-Hand Movement

The object to reach for was a wooden parallelepiped (10 × 2.5 × 5cm, weighting 96g) situated 35 cm away from a starting switch on the table, aligned with participants’ dominant shoulder along the sagittal axis. Each trial (18 trials in total) started with participants holding their fingers in a pinch grip position on the starting switch. A tone signaled that they could start their movement. Participants were instructed to reach, grasp and lift the object at their natural speed, using their dominant hand, then to put the object down and go back to the starting position until the next tone was presented.

#### Forearm Length Estimation

In this task, participants were blindfolded. Each trial (18 trials in total) started with them holding their index finger on the starting switch, waiting for a tone to indicate the start of the estimation. Participants were asked to slide their dominant index finger horizontally (i.e., towards the right/left) on the surface of the table from the starting switch to a final position corresponding to the perceived estimation of their forearm (i.e. perceived distance between wrist and elbow). To avoid estimation bias by visual forearm measurement, the experimenter gave task instructions once the participants were blindfolded. The experimenter also named and touched the elbow and the wrist of each participant while giving the instructions, so that they would know exactly which body part they had to estimate.

#### Tool movement

The tool-use task was composed of four blocks of 12 trials. Each trial started with participants holding the tool “fingers” in a pinch grip position on the starting switch. After a tone indicating the go for the movement, participants had to reach, grasp and lift the same object as previously described at their natural speed, using the tool with their dominant hand. Between each trial, they had to put the object down and go back to their starting position until presentation of the next tone. Each participant underwent a short practice tool-use session at the beginning of the tool session. The tool, based on a commercial grabber (Unger Enterprise Inc, CT, USA), had an ergonomic handle fitted with a lever, a long rigid shaft, and a “hand” with two articulated fingers. It was customized and scaled on children’s height (see Martel et al., 2021 for details). Participants between 123 and 146 cm of height used a 32 cm long tool (DCD group: 4/17; TD group: 5/17). The remaining participants were taller than 147 cm and used a tool of 40 cm long, which is the original length also used for adults (Baccarini et al., 2014; Cardinali et al., 2011, 2009; Martel et al., 2021, 2019), but 100g lighter to prevent fatigue. Participants’ height was collected before the experiment, the lengths of their arm and forearm were measured afterwards.

#### Gesture Imitation proficiency

We quantified participants’ sensorimotor proficiency and compared the performance of DCD and TD groups in a gesture imitation task, initially developed for the assessment of apraxia (De Renzi et al., 1980). Participants were instructed to imitate a set of different gestures performed by the experimenter using their dominant arm/hand (anatomical imitation, not mirrored), while standing. Two training gestures allowed them to familiarize with the task. We also informed them that some gestures would be repeated and emphasized the importance of exact imitation of the gesture (fingers opening, hand orientation etc.). We reminded participants of these instructions several times before the test. The test was composed of 24 gestures: for correct imitation on the first time, participants scored 3 points; 2 points were given if the experimenter had to perform the movement a second time and participants succeeded in imitating it. In case of failure, on the third and last repetition, participants scored 1 for a correct gesture and 0 for inaccurate imitation. We defined the assessment criteria classically used in motor imitation tasks (Rothi et al., 2014). These criteria included the configuration of the arm and/or hand (e.g. extended arm and fist configuration), the limb orientation in space (e.g. palm down) and its target location (e.g. palm on the contralateral shoulder). For sequential gestures, we additionally assessed the correct order and number of occurrences (e.g. three repetitions of fist and hand flat on the table sequence). Any element differing from one of these criteria was considered an incorrect imitation. The maximum score was 72. The same, trained experimenter demonstrated the gestures and evaluated online the imitation for all participants.

The entire procedure lasted about 1h30 including breaks and the signature of the consent forms. DCD participants could undergo the M-ABC assessment on the same day, making the total duration of the session to 3h, or on another day (two sessions of 1h30 each).

### Kinematic recording system

We recorded the spatial localization of the hand and of the tool using infrared light emitting diodes (IREDs) with an Optotrak 3020 (Northern Digital Inc; sampling rate: 200 Hz; 3D resolution: 0.01 mm at 2.25 m distance). Following previous studies using the same paradigm (Baccarini et al., 2014; Cardinali et al., 2012, 2011, 2009; Martel et al., 2021, 2019), we assessed the grasping component of the hand/tool movements by placing IREDs on the thumb and index finger nails of participants’ dominant hand, as well as on the two “fingers” of the tool. The reaching component was evaluated thanks to IRED located on the dominant wrist (styloid process of the radius and distal part of the tool shaft).

For each free-hand movement, we extracted and analyzed off-line several kinematic parameters with a custom-made Matlab program: latencies and amplitudes of wrist acceleration, wrist velocity and wrist deceleration peaks (reaching component), and latency and amplitude of the maximum thumb-index distance (hereafter MGA for Maximum Grip Aperture) (grasping component). We also measured the overall movement time as the time between the beginning of the movement (velocity ≥ 10 mm/s after switch release) and stabilized grasp on the object (before the lift). Extracted parameters were the same for the tool movement task, except that they involved kinematics of the tool instead of the hand. The arm length was estimated using the marker on participants’ index finger by subtracting the starting position to the final one.

### Statistics

We used a Linear Model (LM) on the gesture imitation score of each participant (as it consisted of only one value per participant), and Linear Mixed Models (LMM) on individual trials of each kinematic parameter of tool and free-hand movements, implemented in R (v3.6.1; RStudio v1.1.442; (R Core Team, 2018) with the package lme4 (Bates et al., 2015). Although similar to LM in terms of interpretation, LMM are more adequate for the present study as they take into consideration the variability between participants (Boisgontier and Cheval, 2016), which is especially high in participants with developmental disorders. Inference on main effects and interactions were based on ANOVA Type III Wald chi-square tests (R package car, function Anova; Fox and Weisberg, 2011). We further investigated any significant pairwise difference after post-hoc correction (Tukey method, package emmeans; Russell, 2019).

#### Gesture Imitation task

We performed a LM with factor *Group* (TD/DCD) to assess whether DCD participants had poorer sensorimotor abilities than their TD peers, as expected from the literature.

#### Free-hand movement

We used a LMM with factors *Session* (PRE/POST) and *Group* (TD/DCD) and a random intercept for participants. We were particularly interested in the presence of interactions between the two factors, as they would indicate that tool use modifies free-hand movements of DCD children and early adolescents differently than it does in their TD peers. Additionally, we expected the pattern observed in our subsample of TD participants to conform to the one from the larger sample (Martel et al., 2021), namely that TD children and early adolescents display shorter latencies and higher amplitudes of acceleration, velocity and/or deceleration peaks after tool use.

#### Forearm Length Estimation

A LMM with factors *Session* (PRE/POST) and *Group* (TD/DCD) and a random intercept for participants was used (note that data were missing for one TD and one DCD participant). Interactions between the two factors would indicate that tool use does not affect the conscious forearm representation of DCD children and early adolescents the same way it does for TD participants. Additionally, for the latter, we expected the conscious forearm representation to be reduced after tool use, as previously reported in the larger sample of TD participants (Martel et al., 2021).

#### Tool movement

We used a LMM with factors *Block* (FIRST/SECOND/THIRD/LAST) and *Group* (TD/DCD) and a random intercept for participants. Would the two factors interact, this would show that both groups do not control the tool similarly along blocks. As before, we also anticipated to observe the motor learning effects found in the larger sample of TD participants (Martel et al., 2021), with children and early adolescents displaying shorter latencies and higher amplitudes in the last compared to the first block.

## Results

### DCD participants show poorer gesture imitation abilities than TD participants

The DCD group showed a mean score for gesture imitation of 49.2 ± 8.8. As evidenced by a significant effect of *Group* in the linear model (F(1, 32) = 10.34, p = .003), this was significantly lower than the performance of the TD group who scored at 57.6 ± 6.2 (Figure 2). Noteworthy, when considering the cutoffs from the original test in adults (healthy ≥ 62; apraxia ≤ 53; De Renzi et al., 1980), the two groups did not fall in the same category, underlining their different sensorimotor abilities. Note that this test has been standardized on adults, lower performance are therefore to be expected in the present study (see also Martel et al., 2021), however this can not explain a change in categories between the groups.

**Figure 2.**
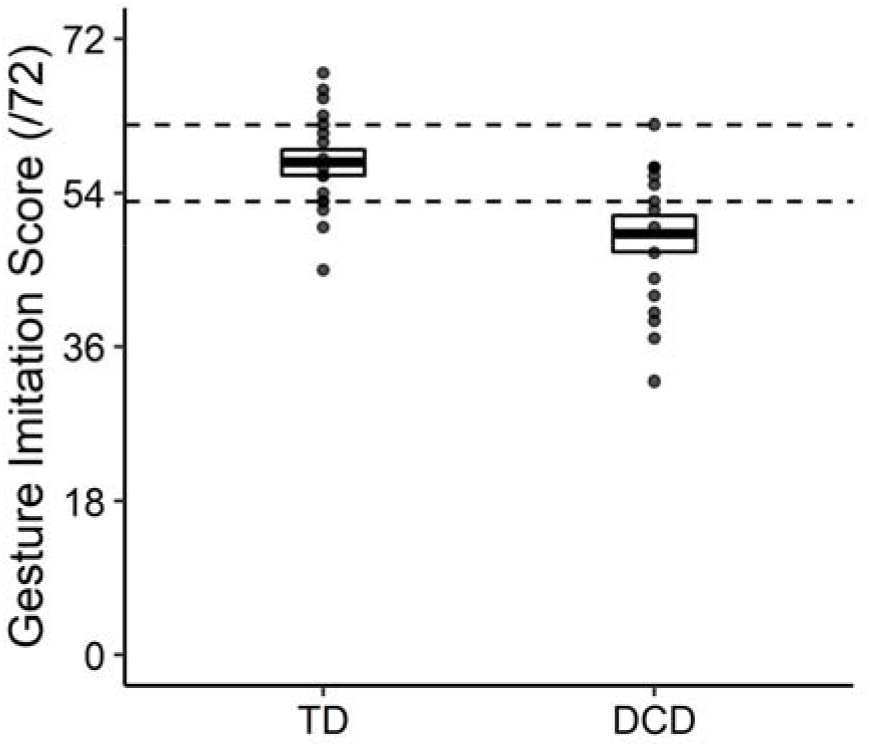
Performance in the gesture imitation test in TD and DCD participants. The thick horizontal lines indicate the mean for each group ± 1 SEM. Single dots represent individual performance. Horizontal dashed lines represent the cutoffs in the original task standardized on adults (healthy ≥ 62; apraxia ≤ 53; De Renzi et al., 1980)

### Tool use affects free-hand movements differently in TD and DCD children and early adolescents

#### Reaching component

The LMM revealed a significant main effect of *Session* on all of the amplitudes (all p < .001) but none of the latencies (all p > .115). Of major interest here, significant interactions between *Session* and *Group* were observed for all the latencies (acceleration: χ^2^(1) = 3.99, p = .046; velocity: χ^2^(1) = 9.04, p = .003; deceleration: χ^2^(1) = 10.9, p < .001). Post-hoc comparisons showed that while the TD participants displayed decreased latencies for two out of three parameters when reaching after tool use (velocity: t = 2.62, p = .044; deceleration: t = 3.46, p = .003; Figure 3), no latency modulation in subsequent free-hand movements was found in the DCD children and early adolescents (velocity: t = −1.63, p = .364; deceleration: t = −1.23, p = .611). Both TD and DCD participants displayed increased amplitudes following tool use, as indicated by the lack of significant *Session* × *Group* interactions (acceleration: χ^2^(1) < .001, p = .928; velocity: χ^2^(1) = .444, p = .505; deceleration: χ^2^(1) = 3.21, p = .073).

**Figure 3.**
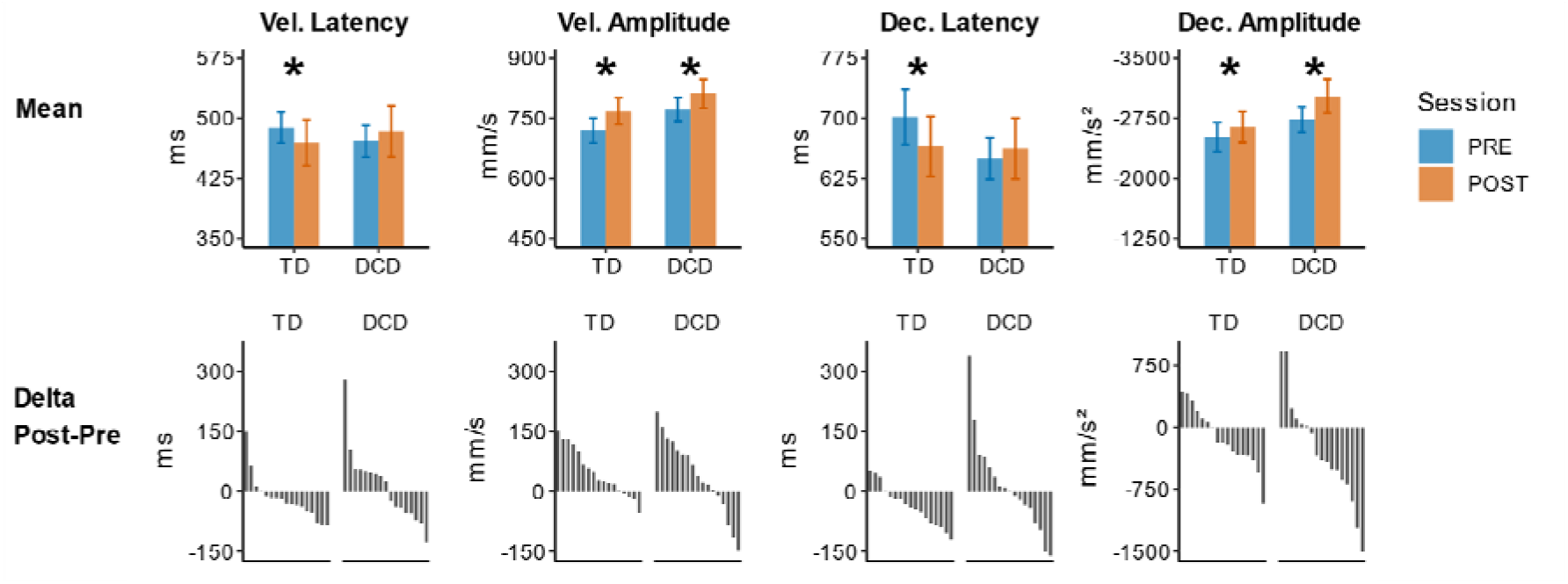
Kinematics of the Reaching component before (blue) and after (orange) tool use (top panel) and individual performance ordered by size (bottom panel) in TD and DCD participants for the latencies and amplitudes of velocity and deceleration. On the bottom panel, positive values indicate that the latency/amplitude increased after tool use, while negative values indicate reduced latency/amplitude. Most of the TD children and early adolescents displayed reduced latencies and increased amplitudes after tool use. DCD children and early adolescents displayed increased amplitudes but no modulation of the latencies. Error bars indicate the means ± 1 SEM. Asterisks denote significance.

#### Grasping component

The factor *Session* significantly affected the latency of MGA (χ^2^(1) = 18.5, p < .001): children and early adolescents opened their fingers earlier after tool use. There was no evidence that DCD participants modulated this parameter differently than their TD peers, with neither a main effect of *Group* (χ^2^(1) = .360, p = .549), nor an interaction between *Session* and *Group* (χ^2^(1) = .683, p = .409). Regarding the MGA, the DCD participants opened their fingers larger than the TD participants with a main effect of *Group* (χ^2^(1) = 16.5, p < .001). There was also a significant main effect of *Session* (χ^2^(1) = 42.7, p < .001) and a significant *Session* × *Group* interaction (χ^2^(1) = 14.2, p < .001). As revealed by post-hoc comparisons, tool use increased subsequent free-hand finger opening in the DCD group (t = −7.12, p < .001) while it did not modulate grip aperture in TD participants (t = − 2.01, p = .186; Figure 4A).

**Figure 4.**
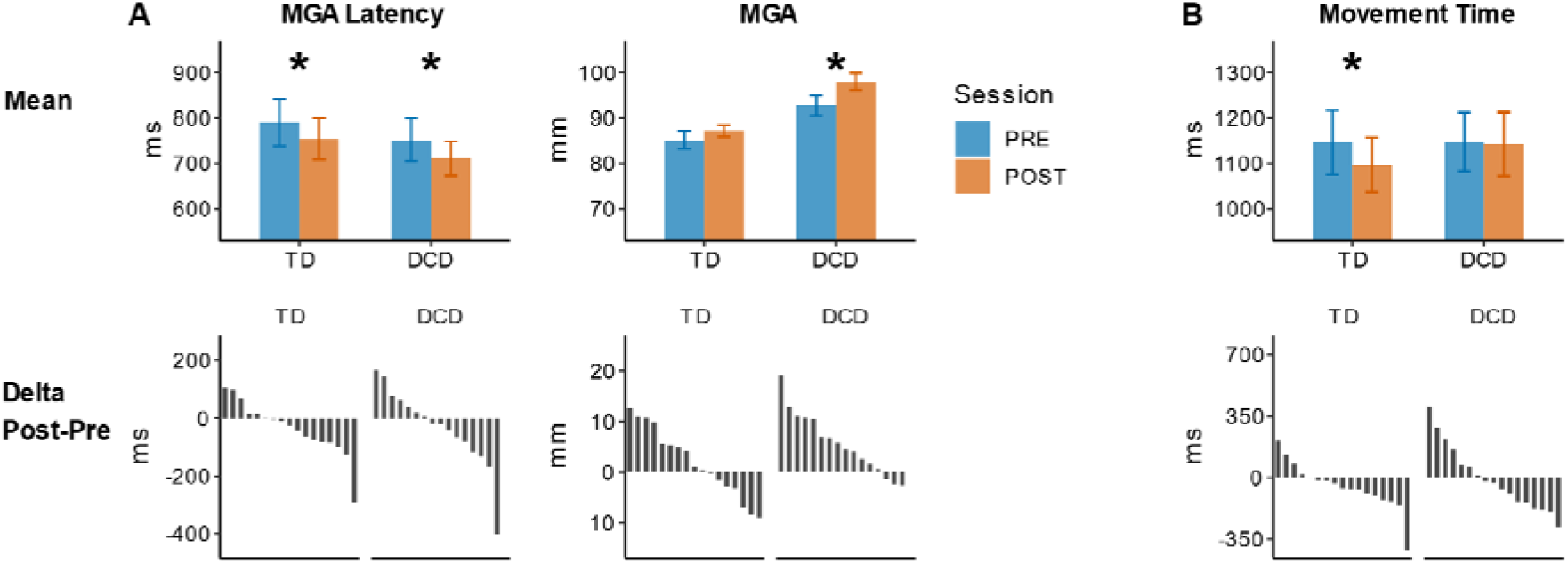
Kinematics of the Grasping component (A) and the Movement Time (B) before (blue) and after (orange) tool use (top panel) and individual performance ordered by size (bottom panel) in TD and DCD participants. On the bottom panel, positive values indicate that the latency/grip aperture and movement time increased after tool use, while negative values indicate reduced latency/grip aperture and shorter movement time. Note that data for the grasping component is missing for one DCD participant. Most of the TD children and early adolescents displayed reduced MGA latencies, which led to reduced movement time after tool use. DCD children and early adolescents opened their fingers earlier and larger after tool use. They however did not display any modulation of their movement time. Error bars indicate the means ± 1 SEM. Asterisks denote significance.

#### Movement Time

The LMM revealed a significant main effect of *Session* (χ^2^(1) = 6.83, p = .009) as well as a significant interaction between *Session* and *Group* (χ^2^(1) = 4.85, p = .028). No significant main effect of *Group* (χ^2^(1) = .073, p = .788) was found. Post-hoc comparisons indicated that free-hand movement time was shorter after tool use in the TD group (t = 3.41, p = .004; Figure 4B), while the duration of the movement was not modulated in the DCD group (t = .291, p = .991).

In sum, the group of TD participants depicted larger amplitudes and reduced latencies during the reaching component of their movement, resulting in shorter movement times. They also displayed a reduced latency, opening their fingers earlier, but no modulation of the amplitude of the grasping component. Instead, DCD children and early adolescents displayed modulations of the amplitudes of the reaching component only, while their latencies were not affected, which did not lead to a change in movement times. They however modulated their grasping component, opening their fingers both earlier and larger after tool use.

### Forearm length estimation is reduced after tool use in both TD and DCD groups

Participants estimated their forearm shorter following tool use, as shown by a significant main effect of *Session* (χ^2^(1) = 55.9, p < .001; Figure 5; mean ± 1 sem: TD: 114% ± 9.2% vs 105% ± 7.8%; DCD: 131% ± 10.3% vs 122% ± 11.2). However, the two groups did not significantly differ in their estimation (no main effect of *Group*: χ^2^(1) = 1.72, p = .189), which was overall slightly overestimated before tool use. There was no evidence that DCD participants modulated their conscious forearm representation differently than their TD peers as shown by the absence of interaction between *Session* and *Group* (χ^2^(1) = .528, p = .467).

**Figure 5.**
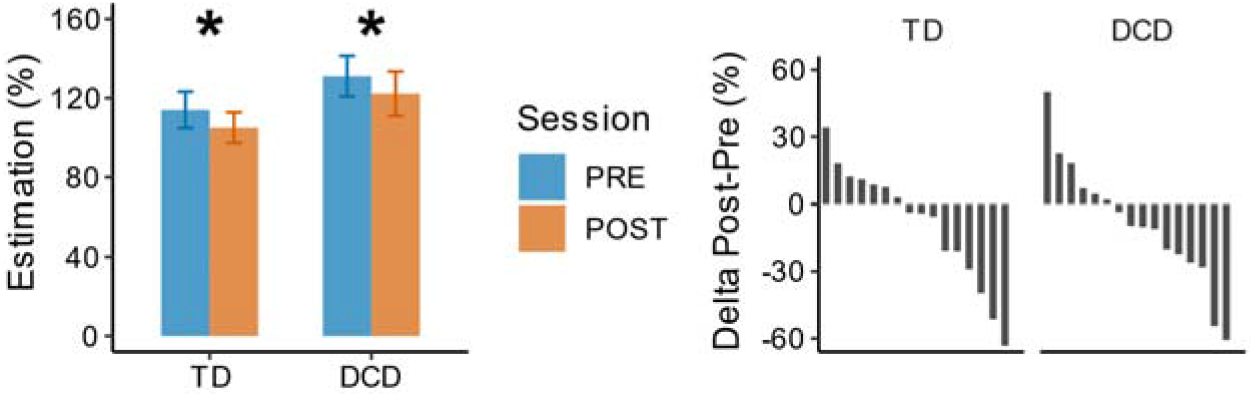
Forearm length estimation (in % of the actual forearm length) before (blue) and after (orange) tool use (left panel) and individual performance ordered by size (right panel). On the right panel, positive values indicate that the perceived forearm length increased after tool use, while negative values indicate reduced forearm length representation. Note that data was missing for one TD and one DCD participant. Most of the TD and DCD children and early adolescents estimated their forearm length to be shorter after tool use. Error bars indicate the means ± 1 SEM. Asterisks denote significance.

### TD and DCD children and early adolescents behave similarly during tool use

#### Reaching component

The LMM revealed a significant main effect of *Block* on all the parameters (all p < .037). Latencies decreased and amplitudes increased gradually from the first to the last blocks of the tool-use session (see Figure 6). Crucially, no evidence indicated that the DCD participants controlled the tool differently to reach for the object than the TD group. The main effect of *Group* was not significant either on latencies (acceleration: χ^2^(1) = .076, p = .783 velocity: χ^2^(1) < .001, p = .988; deceleration: χ^2^(1) = .121, p = .728) or on amplitudes (acceleration: χ^2^(1) = .394, p = .530; velocity: χ^2^(1) = .019, p = .890; deceleration: χ^2^(1) = .329, p = .566). Similarly, there was no significant *Block* × *Group* interaction on most of the latencies (acceleration: χ^2^(3) = 1.60, p = .660; velocity: χ^2^(3) = 3.50, p = .320) and amplitudes (acceleration: χ^2^(3) = .272, p = .965; velocity: χ^2^(3) = 4.15, p = .246; deceleration: χ^2^(3) = 2.72, p = .437). The only significant interaction was observed for the deceleration latency: χ^2^(3) = 8.81, p = .032. Post-hoc comparisons indicated that deceleration latencies were reduced gradually across blocks in DCD participants (1^st^ block differing from the 3^rd^ and 4^th^, p < .001 but not from the 2^nd^: t = 1.46, p = .828; the 2^nd^ differing from the 3^rd^: t = 3.04, p = .049 and the 4^th^: t = 4.88, p < .001). Instead, deceleration latencies did not improve further after the second block in TD participants (the 1^st^ block differing from all the others: all p < .004, in turn not differing between them: all p > .99).

**Figure 6.**
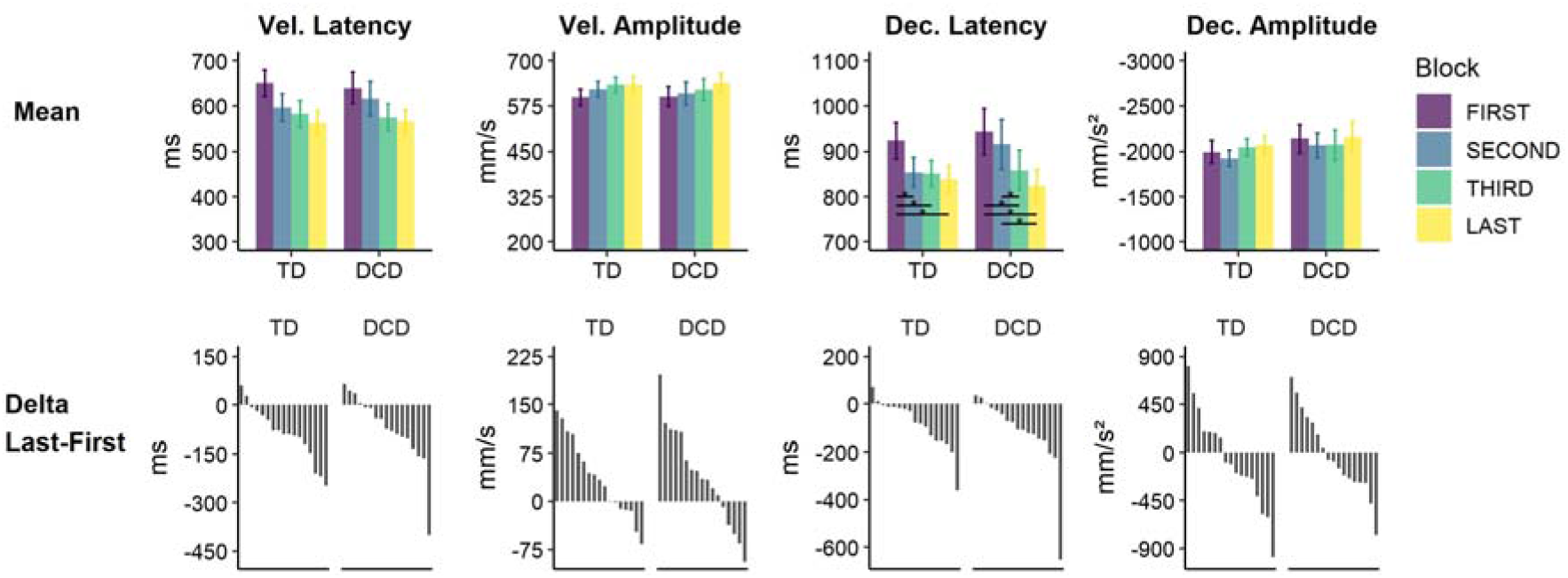
Kinematics of the Reaching component during the four blocks of tool use (top panel) and individual performance (first and last block) ordered by size (bottom panel) in TD and DCD participants for the latencies and amplitudes of velocity and deceleration. On the bottom panel, positive values indicate that the latency/amplitude increased with practice, while negative values indicate reduced latency/amplitude. Most of the TD and DCD children and adolescents displayed reduced latencies and increased amplitudes with tool practice. Error bars indicate the means ± 1 SEM. Asterisks denote significance of the post-hoc comparisons performed because of the significant interaction.

#### Grasping component

Similar to the reaching component, a significant main effect of *Block* (χ^2^(3) = 28.4, p < .001) was found on MGA latency, with participants opening their tool fingers earlier with practice. There was no indication of significant modulation by *Group* (χ^2^(1) = 1.35, p = .245) nor interaction between *Block* and *Group* (χ^2^(3) = 2.06, p = .560). As to the tool maximal grip aperture, the LMM revealed a significant main effect of *Block* (χ^2^(3) = 12.8, p = .005) indicating a larger grip aperture with practice. There was no significant difference between the two groups (χ^2^(1) =. 659, p = .417). The *Block* × *Group* interaction tended to significance (χ^2^(3) = 6.88, p = .076), possibly due to a wider tool grip aperture at the last block compared to the first block for the DCD group.

#### Movement Time

The LMM showed a significant main effect of *Block* (χ^2^(3) = 109, p < .001) and a significant interaction between *Block* and *Group* (χ^2^(3) = 38.0, p < .001). This interaction is likely to be due to a larger modulation between the 2^nd^ and the 4^th^ blocks in DCD compared to TD participants (t = −4.18, p < .001). Post-hoc comparisons further indicated that movement times were longer in the TD group for the first block compared to the other three (all p < .018), which did not differ between them (all p > .988). DCD participants also displayed shorter movement times with practice, but in a more gradual way (1^st^ vs 2^nd^ block: t = .303, p = 1; 2^nd^ vs 3^rd^: t = 3.66, p = .006; 3^rd^ vs 4^th^: t = 5.78, p < .001) (see Figure 7B). Additionally, the significant main effect of *Group* (χ^2^(1) = 4.09, p = .043) showed that DCD participants took more time to reach and grasp the object with the tool than their TD peers, overall tending to display larger time shortening at the end of the session.

**Figure 7.**
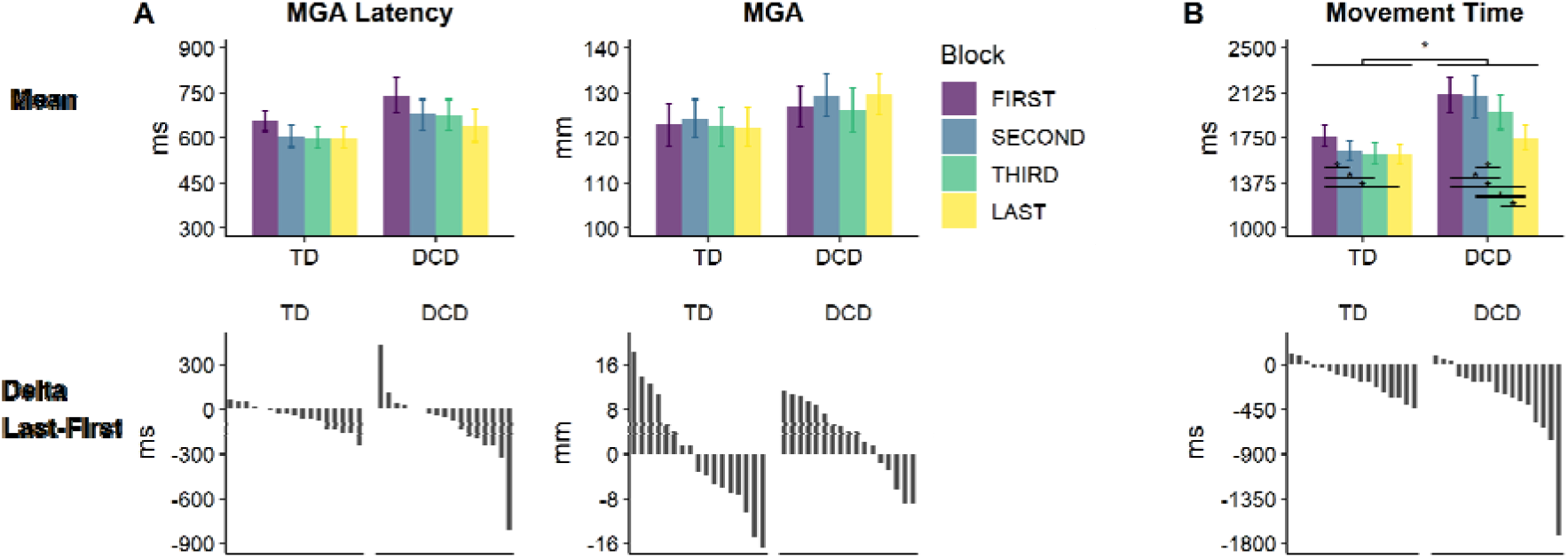
Kinematics of the Grasping component (A) and the Movement Time (B) during the four blocks of tool use (top panel) and individual performance ordered by size (bottom panel) in TD and DCD participants for the first and last blocks. At the bottom, positive values indicate increased latency/grip aperture with practice, while negative values indicate that these parameters decreased. Most of the TD and DCD children and early adolescents displayed reduced latencies, which led to reduced movement time with practice. Additionally, the DCD children and early adolescents opened their tool fingers larger with practice. Error bars indicate the means ± 1 SEM. Asterisks denote significance of the post-hoc comparisons performed because of the significant interaction as well as the main effect of Group.

To summarize, both TD and DCD participants performed faster with the tool with practice and displayed similar modulations of their kinematic parameters across tool-use blocks. Specific interactions however indicated that the shortening of the movement times was stronger in DCD than in TD participants. Additionally, DCD participants displayed a different learning strategy for some parameters. They displayed a gradual reduction of deceleration latencies and movement time across blocks, while TD participants showed stable performance on these parameters by the second block.

## Discussion

In this study, we assessed whether Body Schema plasticity induced by tool-use is affected in children and early adolescents with DCD, as compared to age-, sex- and puberty-matched TD participants. To this aim, we examined tool control in DCD by measuring their motor learning during the use of a mechanical grabber. No major differences between DCD and TD groups were observed, testifying of a skillful tool use in DCD. We then focused on the plasticity of two distinct body representations: the explicit (action free) Body Image (BI) and the implicit (action-related) Body Schema (BS), also referred to as body estimate. Regarding the explicit BI, we compared conscious forearm length estimation of participants before and after tool use. Both groups judged their forearm as being shorter after tool use, as previously reported in a larger cohort of TD participants (Martel et al., 2021) This attests of a preserved explicit body representation plasticity in DCD children and early adolescents. To investigate the implicit BS, we compared participants reach and grasp movements before and after using the tool to reach for the same object. While TD participants showed larger amplitudes and reduced latencies in the kinematic parameters of the reach phase of their movements after tool use, DCD participants displayed modulations limited to the amplitudes of their kinematics. At odds with their preserved Body Image plasticity, the implicit Body Schema plasticity is therefore impaired in DCD. These findings will next be discussed within the internal modelling framework.

### Children and early adolescents with DCD have a preserved ability for motor learning

DCD and TD participants showed no major difference in controlling the tool. With practice, both groups improved in using the tool, in keeping with the findings in a larger sample of TD participants (Martel et al., 2021). This suggests that DCD children and early adolescents were able to use visual and/or proprioceptive feedback and compare them with the predicted one/s. As using a tool is thought to require the building of new sensorimotor associations during development (Ganesh et al., 2014; Martel et al., 2021), it is likely that the predicted feedback were initially imprecise and that participants relied on the more trustable information, that is their actual feedback (Limanowski and Friston, 2020a, 2020b). Their performance suggests they took error signals into account to improve ensuing movements by updating their internal models, to a similar extent their TD peers did. Yet, this is not to imply that TD and DCD participants favored the same sensory modality equally. Indeed, DCD children and early adolescents tended to open their tool larger with practice, in line with what has been previously reported in free-hand movements in DCD children, suggesting that they may particularly rely on vision during movement execution (Biancotto et al., 2011). Noteworthy, the total movement time during tool use in this study was longer and more strongly modulated by practice in DCD than in TD participants. As the end of the movement was defined as the time just before the lift of the object, and the reaching component was comparable across groups, the larger effects induced by tool use on movement time in DCD may emerge from the grasping component. The deceleration phase before object contact is indeed characterized by a low velocity phase when approaching the object to grasp and is known to rely mostly on visual feedback integration. The more gradual modulation of the deceleration latency and movement time in DCD participants further suggests that, even if motor learning is preserved, actions requiring feedback integration might be more problematic in DCD. Both TD and DCD children and early adolescents switched from small opening of the tool and longer movement time at the beginning of the tool-use session, to larger opening at the end, guarantying a safety margin when grasping (Wing et al., 1986) and thus penalizing less the total movement duration; this was more pronounced in the DCD group. The DCD group might be slightly less skilled with the tool, although one would expect differences on the reaching component as well in this case, making this possibility unlikely. Alternatively, a conscious executive strategy could be used by DCD to avoid dropping the object, resulting for instance from reduced self-esteem and perceived competence (Watson and Knott, 2006). Further, as modulation of the grip aperture is particularly related to visual control (e.g. Marino et al., 2010), DCD may have relied more on visual feedback while controlling the tool, while their TD peers could benefit from both their proprioceptive and visual feedbacks. The latter possibility seems more likely as it would agree with studies reporting poorer proprioception in DCD (e.g. Coleman et al., 2001; Li et al., 2015; Smyth and Mason, 1998a, 1997; Smyth, 1994; Visser et al., 1998). It also fits with the impairment observed by the present DCD sample in the sensorimotor imitation task, for which low performance has been linked to diminished proprioception abilities (Assaiante et al., 2014; Cignetti et al., 2013; Martel et al., 2021). Overall, our results in DCD children and early adolescents are in keeping with their preserved abilities for motor learning (Smits-Engelsman et al., 2015), as well as their ability to update inverse models and program adequate motor commands (Gomez and Sirigu, 2015; Smits-Engelsman and Wilson, 2013). As such, DCD might not be limited to a purely motor disorder, but is likely to involve deficits in multi-sensory integration. Most of the DCD children and adolescents in our sample had the same consistent pattern; still they presented co-occurring disorders, in agreement with the 40% prevalence reported in the literature (e.g. Dewey et al., 2002; Flapper and Schoemaker, 2013; Lingam et al., 2010; Loh et al., 2011). Therefore, we cannot exclude that some might have stronger motor impairments than others (see for instance (Cignetti et al., 2018; Lewis et al., 2008). This highlights the necessity for future studies to try correlating experimental findings with several clinical measures.

### Body estimate plasticity is altered in DCD children and early adolescents

As recalled in the introduction, TD children and early adolescents show a tool-induced kinematic pattern opposite to adults (Baccarini et al., 2014; Bahmad et al., 2020; Cardinali et al., 2011, 2009; Martel et al., 2019): after tool use, their kinematics display increased peak amplitudes and decreased peak latencies in several parameters of the reaching phase of their free-hand movements, resulting in shorter movement times (Martel et al., 2021). While it is presently uncertain whether it results from vision-biased motor control of tools, or the establishment of new sensorimotor associations (or both), this pattern is consistent with a change in the arm length estimate, in the direction of shortening. Indeed, previous work in TD children and early adolescents indicates that their body estimate is plastic and can be updated within the internal models along a trajectory that goes first in the direction of a shorter arm representation, to then attain the adult pattern (a longer representation) several years later (Martel et al., 2021).

Unlike their TD peers, DCD children and early adolescents only displayed increased peak amplitudes after tool use, no decrease in peak latencies being observed. In other words, they did not reach the object as if their arm was shorter after tool use. This is particularly interesting because the partial modulation of their kinematics was present despite their motor control of the tool was comparable to their matched TD peers, clearly suggesting that body estimate (BS) plasticity is altered in DCD. Selective modulation of the amplitudes but not the latencies in the reaching kinematics has been previously observed after tool-use imagery (Baccarini et al., 2014). In that study, healthy adults displayed the typical kinematic signature of body representation plasticity after having imagined using a tool, but only on their movement amplitudes. Thus, one might think that altered updating plasticity in DCD might emerge as a consequence of tool imagery, that is participants would rely more on their predicted feedback than their received ones during tool use. DCD children and early adolescents do have the ability to engage in motor imagery (for review, Adams et al., 2014; Barhoun et al., 2019; Gabbard and Bobbio, 2011), indicating that they are able, to some extent, to predict the consequences of their motor commands. However, motor learning during tool use was comparable across TD and DCD groups, suggesting that DCD participants did not solely rely on tool imagery during tool use, even though they might have attributed a different weight to the sensory feedback than their peers.

Interestingly, previous work in adults also showed significantly larger modulations of the amplitudes after tool use when vision and proprioception were both present, compared to when only proprioception was available (Martel et al., 2019). It is thus tempting to speculate that vision might preferentially affect amplitudes of free-hand movements after tool use, in agreement with the notion that DCD favor vision over proprioception when both are available (e.g. Biancotto et al., 2011; Smyth and Mason, 1998b). This is further in line with their difficulties in the sensorimotor imitation task reported here and, more generally, with previous reports of poorer proprioception in DCD as compared to TD children (e.g. Li et al., 2015; Visser et al., 1998). While future studies are needed to specifically address the role of vision and proprioception in altering DCD’s body estimate processes, the present findings also open the possibility that such altered BS plasticity may in turn affect the proper internal modelling. This study thus brings evidence in favor of an alternative theoretical account of the DCD etiology. Our findings point to a deficit in the plasticity of the body representation used to plan and execute movements. Though not mutually exclusive, this widens the theoretical perspective under which DCD should be considered: DCD may not be limited to a problem affecting the internal models and their motor functions, but may concern the state of the effector they have to use. Further studies are also needed to determine whether these behavioral impairments relate to abnormal brain activation previously reported in DCD, such as in the somatosensory and motor areas (e.g. Gomez and Sirigu, 2015; Kashiwagi et al., 2009; Zwicker et al., 2010).

### Children and early adolescents with DCD have a preserved ability to update the explicit metrics of their body

In healthy adults, the explicit representation of the body metrics (or Body Image) has been shown to be largely immune to the effects of tool use. When required to indicate their forearm length using the same paradigm as in the present study, adults’ performance shows that the explicit knowledge of their forearm length is not modified after tool use (Bahmad et al., 2020; Cardinali et al., 2012, 2011; Martel et al., 2019). Yet, this is not the case during development: TD children and adolescents judge their forearm as being shorter after tool use (Martel et al., 2021), a sign of their Body Image plasticity. Here, this pattern was observed not only in TD but also in DCD participants, highlighting their ability to access a conscious metrics of their limb length, as well as to update it after tool use. Interestingly, if DCD participants predominantly used visual feedback during tool use, and possibly to a larger extent than their TD peers, this may have led to an update of the explicit representation in the direction of a shorter limb, without any update of the implicit body estimate. For the first time to our knowledge, we report a dissociation between access to the implicit body estimate for action (Body Schema) and the conscious Body Image in DCD: while the former appears impaired, the latter is preserved. In this respect, some perceptual problems have been attributed to dys-functioning of the dorsal stream in several developmental disorders (Pisella et al., 2019), with parietal but not occipital visuo-spatial dysfunction being particularly involved in DCD (Nobusako et al., 2018; Pisella et al., 2020, 2019). This could indicate that plasticity of the Body Image mostly involves visual information processed in the occipital part of the dorsal stream, which may be intact in DCD. The parietal part of the dorsal stream, involved in the plasticity of the implicit body estimate, might instead be impaired, in keeping with the proposed role of state estimator for the posterior parietal cortex (Medendorp and Heed, 2019; Shadmehr and Krakauer, 2008), and parietal dysfunction in DCD (Debrabant et al., 2016; Kashiwagi et al., 2009; Zwicker et al., 2011).

### Clinical implications

Although somewhat neglected in research (Gomez and Sirigu, 2015), body representations are central in the clinical remediation of DCD. Physical therapists usually focus their approach on the body rather than motor disorders. They work with children to improve their body awareness for instance and find compensatory strategies. The clinical assessment of the ability to point or name several body-parts is generally preserved in DCD children, though its potential relationship with their ability to access the conscious metrics of their arm, also preserved, needs to be elucidated in further studies. On the contrary, they experience difficulties in adjusting their body when their posture is challenged, or when they are asked to run fast or jump during clinical assessment; these situations require them to access their body representation for action. We believe that research should learn from field observations, and that the clinical community would in turn benefit from systematic evaluation of body representations and their plasticity in DCD research. A better understanding of the role of body representations in internal modelling and of how the body interacts with sensory information will eventually help remediation in DCD children and possibly adults, for whom knowledge of the deficits is still partial, making more difficult to find efficient remediation approaches.

## Conclusion

This study reveals that the update of the body estimate used for motor control is altered in children and early adolescents with DCD. Their preserved motor learning, together with clues of preponderant reliance on vision to control hands and tools, also point towards a body-related deficit. Altogether, our findings suggest that children and early adolescents with DCD have trouble when comparing their predicted and received feedback, leading to difficulties in their body estimate. Although DCD has long been considered a motor disorder, our work emphasizes the need to more thoroughly investigate their body representations, as this may also be useful to take into account for benefiting future remediation strategies.

## Acknowledgements

This work was supported by the ANR Samenta (ASD-BARN 01502) and the ANR-16-CE28-0015 Developmental Tool Mastery to AR & AF. It was performed within the framework of the LABEX CORTEX (ANR-11-LABX-0042) of Université de Lyon. MM was supported by a grant from the French Ministry of Higher Education and Research. We thank all children and teenagers who participated in this study and their parents. We thank S Alouche, JL Borach, C Fressard, R Makine, and S Terrones for administrative and IT support and F. Volland for customizing the tools. We thank M.-C. Thiollier and M. Vernet from the Health Network Dys/10 as well as the associations 123 Dys and DFD01 for their help in recruiting the DCD population. We thank V. Veillith d’Aubarede for her help in recruiting the DCD population and the M-ABC assessments, as well as her insights on the clinical aspects of DCD.

## Data availability statement

All data and code used in this study are available on the website of the Open Science Framework (https://osf.io/spxdz/).

